# A pro-oxidant combination of resveratrol and copper down-regulates multiple biological hallmarks of ageing and neurodegeneration

**DOI:** 10.1101/2022.05.04.489500

**Authors:** Kavita Pal, Gorantla V Raghuram, Jenevieve Dsouza, Sushma Shinde, Vishalkumar Jadhav, Alfina Shaikh, Bhagyeshri Rane, Harshali Tandel, Dipali Kondhalkar, Shahid Chaudhary, Indraneel Mittra

## Abstract

Several hundred billion to a trillion cells die in the body every day, and cell-free chromatin particles (cfChPs) that are released from them enter into the extracellular compartments of the body, including into the circulation. We have earlier reported that cfChPs can readily enter into healthy cells to damage their DNA, activate apoptotic pathways and induce inflammatory cytokines. We hypothesized that repeated lifelong assault on healthy cells by cfChPs is the underlying cause of ageing, and that the ageing process could be retarded by deactivating cfChPs. The latter can be effected by oxygen radicals that are generated upon admixing the nutraceuticals resveratrol (R) and copper (Cu). Using confocal microscopy and antibodies against DNA and histone we detected copious presence of extra-cellular cfChPs in brain of ageing mice, and observed that these were deactivated / eradicated following prolong oral administration of small quantities of R-Cu. Deactivation / eradication of cfChPs was associated with down-regulation of several biological hallmarks of ageing in brain cells which included reduction in: 1) telomere attrition, 2) amyloid deposition, 3) DNA damage, 4) apoptosis, 5) inflammation, 6) senescence, 7) aneuploidy and 8) mitochondrial dysfunction. At a systemic level, R- Cu treatment led to significant reduction in blood levels of glucose, cholesterol and C-reactive protein. These results suggest that cfChPs may be global instigators of ageing and neurodegeneration, and that therapeutic use of R-Cu may help to retard the process of ageing.

## Introduction

With progressively increasing longevity, the human race is facing a parallel increase in ageing related degenerative disorders which can severely compromise quality of life. It is predicted that, globally, the number of people age 60 years or above will grow by 38%, from 1 billion to 1.4 billion, outnumbering the youth during the next ten years [1]. The United Nations General Assembly has declared 2021–2030 the Decade of Healthy Ageing, with the ultimate goal to find therapeutic interventions which will simultaneously delay the many conditions associated with ageing [1, 2]. It is argued that healthy ageing should be considered as the ultimate preventive medicine [3]. Ageing is characterised by a myriad pathological processes which lead to gradual deterioration of structure and function of all cells and tissues of the body [4], and is associated with a multitude of degenerative disorders such as Alzheimer’s disease [5], cardiovascular diseases [6], diabetes [7], and cancer [8]. Although many theories of ageing have been advanced [9, 10], none can comprehensively explain the numerous changes that accompany this multidimensional process.

DNA damage and chronic inflammation are two cardinal features of ageing [11, 12]. In this context, we have reported that cell-free chromatin particles (cfChPs) that are released from the billions of cells that die in the body every day, and enter into the extracellular compartments of the body, can be readily internalised by healthy cells wherein they inflict dsDNA breaks, activate apoptotic pathways and induce inflammatory cytokines [13, 14]. This has led us to hypothesise that repeated lifelong assault on healthy cells by cfChPs may be the underlying cause of ageing [15, 16]. Our group has successfully isolated and characterised cfChPs from human serum, which upon EM examination revealed extensive size heterogeneity ranging between ∼10nm and ∼1000nm [13]. We have also reported that blood levels of cfChPs increase with age [17].

Our pre-clinical studies have led to the identification of a novel pro-oxidant combination of the nutraceuticals resveratrol (R) and copper (Cu) which deactivates cfChPs via the medium of oxygen radicals [18–20]. R is a well-known anti-oxidant which has been extensively researched for its health benefits [21]. However, and surprisingly, it acts as a pro-oxidant in presence of Cu, which is also a widely researched nutraceutical [22]. That oxygen radicals are generated when R and Cu are admixed was first demonstrated by Fukuhara *et al* [23]. They showed that R acts as a catalyst to reduce Cu (II) to Cu (I) resulting in generation of oxygen radicals which cleaved plasmid pBR322 DNA [24]. We have extended these findings to show that a combination of R and Cu can degrade genomic DNA and RNA [25], and can deactivate cfChPs *in vivo* by degrading their DNA component [18–20, 25]. We have further observed that, paradoxically, the DNA degrading activity of R-Cu increases as the molar concentration of Cu is gradually reduced with respect to R [25]. On the basis of this finding, in the present study, we kept the molar ratio of R : Cu at 1:10^-4^.

We have reported that a combination of R and Cu, when used at a ratio of 1:10^-4^, has therapeutic effects in several pre-clinical conditions associated with elevated extracellular levels of cfChPs [18–20]. For example, orally administered R-Cu can ameliorate toxic side effects of chemotherapy [18], and radiation therapy [19], and prevent bacterial endotoxin induced cytokine storm and fatality in mice [20]. Our early results also suggest that R-Cu is therapeutically effective in humans. An observational study showed that orally administered R-Cu to patients with severe Covid-19 led to reduction in mortality by nearly 50% [26]. We have also reported that grade III-IV mucositis could be significantly reduced by orally administered R-Cu in patients receiving high dose chemotherapy and bone marrow transplant for multiple myeloma [27]. R-Cu treatment also led to significant reduction in blood levels of inflammatory cytokines in that study.

Oxygen radicals that are generated upon oral administration of R-Cu are readily absorbed from the stomach to have systemic effects in the form of deactivation / eradication of extracellular cfChPs [18-20, 26, 27]. In the present study, we have taken advantage of cfChPs deactivating property of R-Cu to investigate whether prolonged administration of R-C to ageing mice will retard the hallmarks of ageing and neurodegeneration. The dose of R used in our study was 1mg/Kg, and that of Cu was 0.1μg/Kg, given by oral gavage twice daily. This dose of Cu was 20,000 times less, and that of R 5 times less, than those that have been used in pre-clinical studies to investigate their health promoting properties by other investigators [28, 29].

Using confocal microscopy and antibodies against DNA and histone we detected copious presence of extra-cellular cfChPs in brain of ageing mice, and observed that cfChPs were deactivated / eradicated following prolonged oral administration of R- Cu. Deactivation / eradication of cfChPs was associated with down-regulation of multiple biological hallmarks of ageing in brain cells. At a systemic level, R-Cu treatment led to significant reduction in blood levels of glucose, cholesterol and C- reactive protein. Taken together, our results suggest that cfChPs act as global instigators of ageing and neurodegeneration, and that therapeutic use of R-Cu may help to make healthy ageing an attainable goal.

## Methods

### Animal Ethics Approval

The experimental protocol of this study was approved by the IAEC under permission No.16/2015. The experiments were carried out in compliance to the IAEC animal safety and ARRIVE guidelines.

Institutional Animal Ethics Committee (IAEC) of Advanced Centre for Treatment, Research and Education in Cancer, Tata Memorial Centre, Navi Mumbai, India maintains respectful treatment, care and use of animals in scientific research. It aims that the use of animals in research contributes to the advancement of knowledge following the ethical and scientific necessity. All scientists and technicians involved in this study have undertaken training in ethical handling and management of animals under supervision of FELASA certified attending veterinarian.

### Source of resveratrol and copper

The sources of R and Cu were: Resveratrol (Trade name—TransMaxTR, Biotivia LLC, USA (https://www.biotivia.com/product/transmax/); Copper (Trade name— Chelated Copper, J.R. Carlson Laboratories Inc. USA (https://carlsonlabs.com/chelated-copper/).

### Animals and R-Cu dosing

Inbred C57Bl/6 mice obtained from the Institutional Animal Facility were maintained following by our Institutional Animal Ethics Committee standards. They were housed in pathogen-free cages containing husk bedding under 12-h light/dark cycle with free access to water and food. The HVAC system was used to provide controlled room temperature, humidity and air pressure.

The study comprised of 24 C57Bl/6 mice, 12 of which were male and 12 were female. Four mice of either sex were sacrificed when they were 3 months old, and acted as young controls. The remaining 16 mice (8 male and 8 female) were allowed to age until they were 10 months old and divided into two groups: 1) Ageing control mice (N=4 of each sex), and 2) R-Cu treated ageing mice (N=4 of each sex). Animals of both groups were sacrificed after 12 months when they were 22 months old.

R-Cu was administered twice daily by oral gavage for 12 months (from 10 months to 22 months) at a dose of 1mg/Kg of R and 0.1μg/kg of Cu. R, being insoluble in water, was administered as water suspension (100μL), and Cu was administered as a water-based solution (100μL). The ageing control mice were given saline (100μL) twice daily by oral gavage. Our previous studies had shown this dose of R-Cu to be effective in multiple other pre-clinical conditions [18–20].

Reduced physical activity and weight loss of mice were used as humane endpoints of study. At appropriate time points mentioned above, blood was collected via retro- orbital route under isoflurane anaesthesia for serum separation. Animals were then sacrificed under CO2 atmosphere by cervical dislocation under supervision of FELASA trained animal facility personnel. After sacrifice, brain was harvested from all animals, fixed in 10% formalin and paraffin blocks were prepared for further analysis.

### Reagents, antibodies and kits

Details of commercial sources and catalogue numbers of reagents, antibodies and analytical kits used in this study are given in supplementary table 1.

### Assessment of superoxide Dismutase (SOD) levels in brains cells

Expression of SOD in brain cells was estimated using standard immunofluorescence (IF) technique as described by us earlier [18, 20].

### Detection of cfChPs in extra-cellular spaces of brain by fluorescence immune- staining and confocal microscopy

Immune-staining for DNA and histone H4 followed by confocal microscopy was performed on FFPE sections of brain as described in detail by us earlier [20]. Fluorescence intensity of five randomly chosen confocal fields (∼ 50 cells in each field) was recorded, and mean fluorescence intensity (MFI) (± S.E.M) was estimated.

### Assessment of telomere abnormalities

#### Telomere length estimation by qRT-PCR

The average telomere length from the brain tissue was estimated using a highly sensitive quantitative real-time PCR (qRT-PCR) technique [30, 31]. Genomic DNA was isolated from brain tissue using DNeasy blood & Tissue Kit (Qiagen, Hilden, Germany). DNA quantification was performed using a Nanodrop^TM^ spectrophotometer (Thermo Fisher Scientific, Waltham, USA). Ten nanogram of DNA was used in 5 µl of 1 x SYBR Select Master Mix (Applied Biosystems, Foster City, CA, USA), 250 nM of both telomere specific primers or 350 nM of 36B4 primers to a total volume of 10 µl reactions. The thermal cycling conditions for both Telomere and 36B4 are: initial denaturation of 95°C for 10 min followed by 40 cycles of 95 °C for 15 sec, 60 °C for 30 sec & 72 °C for 30 sec. The sequence for telomere-specific forward and reverse primers (Sigma-Aldrich) are 5’ CGG TTT GTT TGG GTT TGG GTT TGG GTT TGG GTT TGG GTT 3’ & 5’ GGC TTG CCT TAC CCT TAC CCT TAC CCT TAC CCT TAC CCT 3’. The sequences for acidic ribosomal phosphoprotein (36B4) – specific forward and reverse primers are 5’ ACT GGT CTA GGA CCC GAG AAG 3’ and 5′TCA ATG GTG CCT CTG GAG ATT 3’, respectively).

Genomic DNA isolated from the spleen of an individual mouse was used as a reference DNA and serially diluted for telomere and 36B4 PCR. All samples were assayed in duplicate on a QuantStudio^TM^ 12K Flex Real-Time PCR System (ThermoFisher) using 384-well block. Standard curves were generated and the relative input amount for both telomere and 36B4 were calculated. The average of the ratio of telomere and 36B4 was reported as the average telomere length.

#### Telomere Q-FISH

Quantitative telomere FISH was performed using Cy3-labeled peptide nucleic acid (PNA) telomere probes (supplementary table 1). FFPE sections of brain were de- paraffinized and serially dehydrated in absolute ethanol series (70/80/100 %) followed by antigen retrieval in sodium citrate buffer (pH 6) at 90°C in water bath and cooled to room temperature. Sections were dehydrated in alcohol series and denatured at 75°C for 6 minutes. Sections were then hybridised with PNA telomere probes overnight at 37 °C. Unbound probes were washed with 2X saline sodium citrate (SSC) buffer followed by 4X SSC at 56 °C for 3 minutes each. The sections were finally washed in 4X saline sodium citrate Tween-20 (SSCT) buffer at RT and mounted in VectaShield DAPI. Images were acquired and analysed using Applied Spectral Bio-imaging system (Applied Spectral Imaging, Israel). Images of ∼500 interphase nuclei were acquired using SpotScan Software 8.1 (Applied Spectral Imaging, Israel) in a multichannel 3-D mode with a constant exposure of 1000 milliseconds for Cy3 (telomeres) and 150 milliseconds for DAPI (nuclei) throughout the experiments. Each 3D image comprised of a stack of 11 focal planes per cell with a sampling distance of 500 nm along the *z* direction and 107 nm in the *xy* direction. Images were de-convoluted and telomere numbers per nucleus was estimated using Spot Count algorithm in SpotView software 8.1. Number of telomere aggregates per nucleus was estimated visually.

### Assessment of **β**- amyloid deposition in brain and BDNF in serum

Detection of amyloid deposition in brain was examined by confocal microscopy on FFPE sections following fluorescent-immune staining using primary antibody against β-amyloid and an appropriate secondary antibody (supplementary table 1). Serum BDNF was estimated by ELISA using a commercial kit (supplementary table 1) according to manufacturer’s instructions.

### Assessment of DNA damage, apoptosis and Inflammation in brain cells

γ-H2AX, active caspase-3 and NF-kB expression were analysed on FFPE sections of brain tissue by standard IF method using appropriate antibodies (supplementary table 1) as described by us earlier [18, 20].

### Assessment of senescence in brain cells

Assessment of biomarkers of senescence was performed on FFPE sections of brain tissue which included: 1) co-localisation of telomere and γ-H2AX IF signals using Immuno-FISH; 2) co-localization of 53BP1 and pro-myelocytic leukemia-nuclear bodies (PML-NBs) IF signals; 3) p16^INK4a^ expression by IF using appropriate antibodies.

### Assessment of aneuploidy in brain cells

Aneuploidy was assessed with respect to chromosome numbers 7 and 16 by FISH using chromosome specific probes on FFPE sections of brain tissues. Sections were de-paraffinised and dehydrated in alcohol series (70-100 %) followed by antigen retrieval in sodium citrate buffer (pH 6) at 90°C in water bath and then cooled to room temperature. Sections were hybridised with chromosome 7 and 16 specific DNA probes overnight at 37°C. Unbound probes were washed off with 2X SSC followed by 0.4X SSC at 70°C for 3 minutes each. The sections were finally washed in 4X SSCT and mounted in VectaShield DAPI. Images were acquired and analysed using Applied Spectral Bio-imaging system (Applied Spectral Imaging, Israel). Number of fluorescent signals per nucleus was counted, and signals more or less than 2N were considered as evidence of aneuploidy. Five fields containing ∼ 500 nuclei were analysed and average number of signals per nucleus was calculated.

### Assessment of mitochondrial dysfunction in brain cells

Mitochondrial dysfunction was analysed on FFPE sections of brain tissue on the basis of expression of the mitochondrial trans-membrane protein TOM20 by IF. Estimation of volumetric changes in TOM20 expression was performed using IMARIS software (Bitplane Technologies, US). Mean volume (in x–y–z planes) was calculated for 5 images (>2,000 mitochondria) for each brain section.

### Assessment of systemic metabolic dysfunction

Serum glucose and cholesterol levels were estimated using an automated instrument (Dimension EXL with LM, Siemens). Serum C-reactive protein (CRP) levels were measured using a commercial ELISA kit (supplementary table 1) as per the manufacturer’s protocol.

### Statistical analysis

Statistical analyses were performed using GraphPad Prism 6 (GraphPad Software, Inc., USA. Version 6.0). Mean (± SEM) values for four mice in each group for both sexes were compared using non-parametric unpaired student’s t test, separately for both sexes.

## Results

### R-Cu up-regulates SOD in brain cells

As a first step, we investigated whether oral R-Cu treatment might have led to generation of free-radicals in the brain. As expected, ageing mice showed significant reduction in SOD levels in brain cells (p<0.05 and p<0.01 in female and male mice, respectively) (Fig 1). However, R-Cu treatment led to marked increase in SOD levels that were similar to that detected in young control mice (p<0.01 in both female and male mice) (Fig 1). This finding suggested that oxygen radicals were being generated *in vivo* following R-Cu treatment, and that they had entered into brains cells. The latter in an attempt to eliminate the invading oxygen radicals had activated an anti-oxidant defence mechanism by up-regulating the anti-oxidant enzyme SOD.

**Figure 1.**
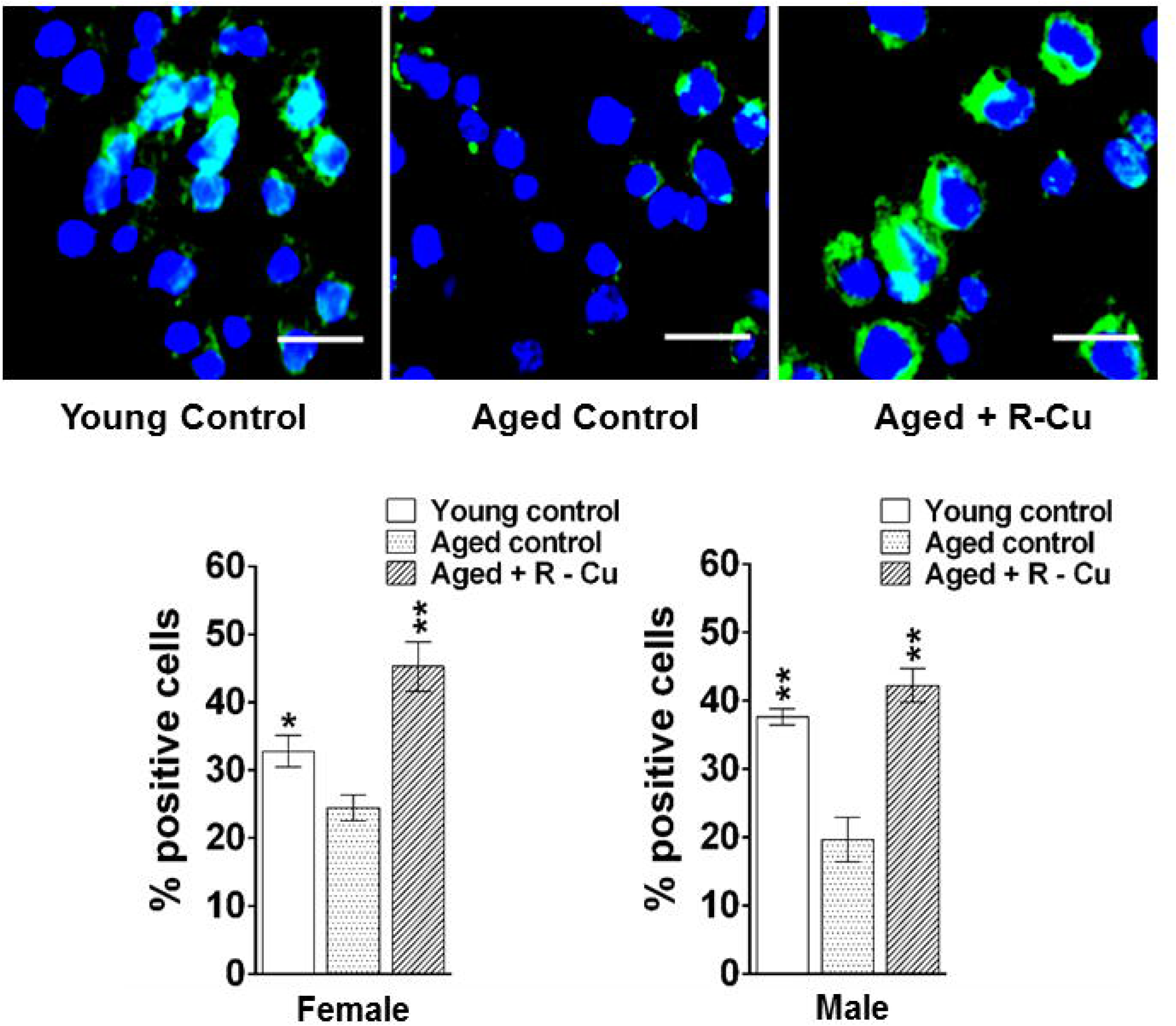
Loss of SOD activity in brain cells of ageing mice and its reversal by R-Cu treatment. Representative IF images of SOD expression in brain cells (upper panel) (Scale bar = 10 µm), and quantification of SOD levels expressed as histograms (lower panel). For each slide 1000 cells were analysed and percent cells positive for SOD were calculated. Bars represent mean ± SEM values. N=4 animals in each group of both sexes. Values of young control mice and R-Cu treated ageing mice were compared with those of ageing control mice, and statistical analysis was performed by two-tailed Student’s t test. * p< 0.05; ** p < 0.01.

### Detection of copious presence of cfChPs into extra- cellular spaces of ageing brain and their deactivation / eradication by R-Cu

Confocal microscopy of FFPE sections of ageing mouse brain was performed following fluorescent immuno-staining with anti-DNA (red) and anti-histone (green) antibodies. Upon superimposing DNA and histone fluorescent images, copious presence of cfChPs (yellow fluorescent signals) were detected in the extracellular spaces of brain of ageing mice (Fig 2). cfChPs were virtually eliminated following one year R-Cu treatment. This finding suggested, in support of our hypothesis, that oxygen radicals generated in the mouse brain had effectively deactivated / eradicated the profusion of cfChPs that are present in the extra-cellular spaces of ageing mouse brain. It should be noted that not all red (DNA) and green (histone) fluorescent signals strictly co-localise. This may have resulted from unevenness of cut surfaces of FFPE sections which prevented the respective antibodies to access the DNA and Histone epitopes on cfChPs. Quantification of MFI of yellow fluorescent cfChPs signals showed a remarkable reduction in cfChPs in extracellular spaces of ageing brain following one year treatment with R-Cu (p< 0.01 in both sexes) (Fig 2, right hand panel).

**Figure 2.**
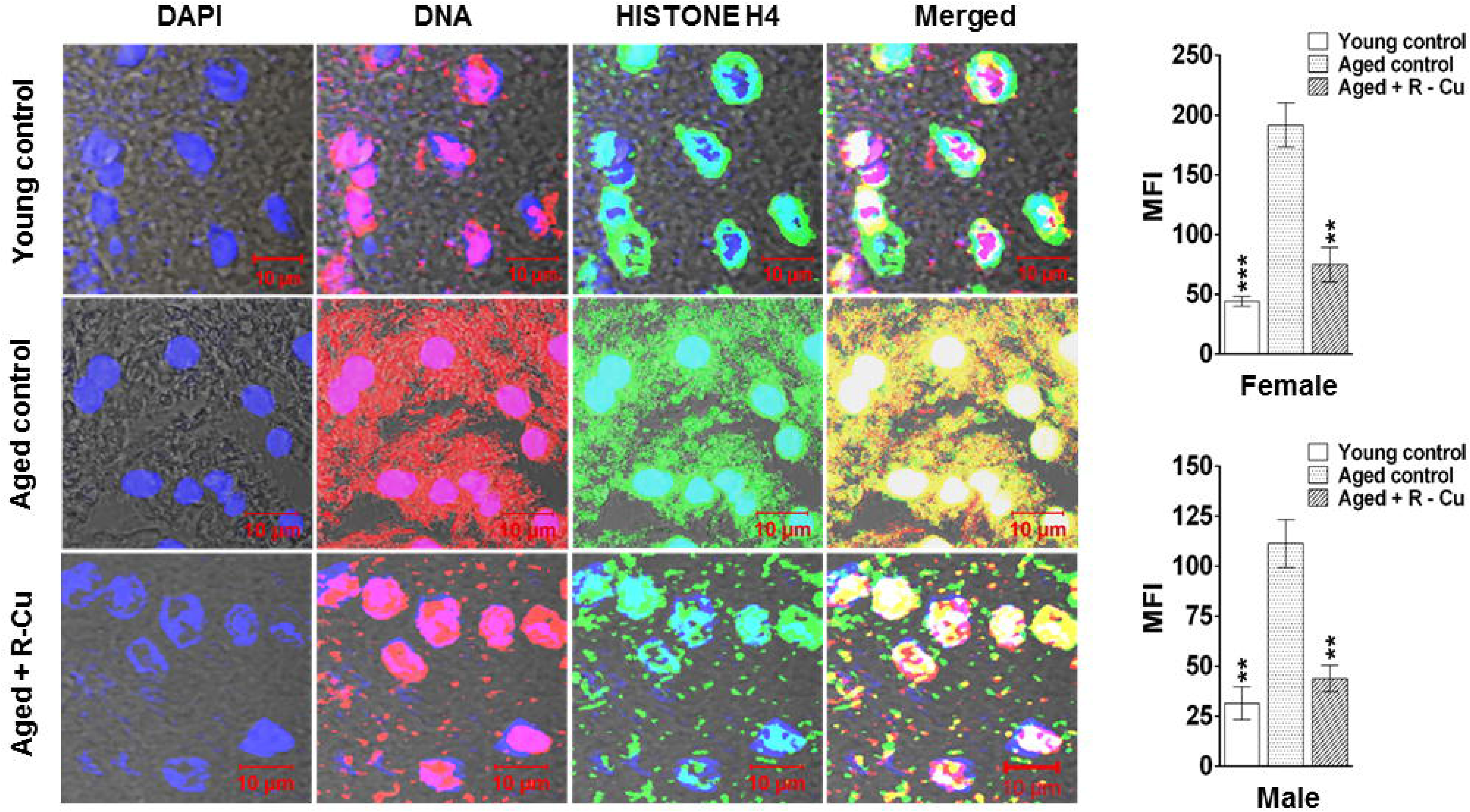
Copious presence of cfChPs into extra- cellular spaces of ageing brain and their deactivation / eradication by R-Cu treatment. Representative confocal images of FFPE sections following fluorescent immuno-staining with anti- DNA (Red) and anti-histone antibodies, showing co-localization of red and green signals to generate yellow / white coloured particles which represent cfChPs (left hand panel). Quantification of yellow IF signals expressed as histograms (right hand panel). Fluorescence intensity of five randomly chosen confocal fields (∼ 50 cells in each field) from each section was recorded. Bars represent mean ± SEM values. N=4 animals in each group of both sexes. Values of young control mice and R-Cu treated ageing mice were compared with those of ageing control mice, and statistical analysis was performed by two-tailed Student’s t test. ** p < 0.01, *** p < 0.001.

### R-Cu prevents telomere abnormalities in ageing brain cells

Telomeres play a central role in cellular changes associated with ageing [32]. Telomere attrition, telomere loss and telomere aggregation are cardinal features of ageing [32, 33]. We estimated telomere length in brain cells by qRT-PCR and observed a significant reduction in telomere length in mice of both sexes (p< 0.0001 and p< 0.01 in female and male mice, respectively) (Fig 3A). However, R-Cu treatment restored telomere length to a significant degree only in female mice (p<0.001), but not in male mice (Fig 3A). With respect to number of telomere signals per brain cell, a reduction was seen in ageing mice of both sexes (p<0.01 in both sexes), which was again significantly restored following R-Cu treatment in female (p<0.01) but not in male mice (Fig. 3B & 3C). A similar observation was made with respect to aggregation of telomeres, which was significantly increased in ageing mice of both sexes (p<0.001 and p<0.01 in female and male mice, respectively), but was significantly reduced following R-Cu treatment only in female (p<0.01), but not in male mice (Fig. 3B & 3D). Thus, overall, with respect to telomere abnormalities, R- Cu was found to be effective in restoring telomere abnormalities in female but not in male mice.

**Figure 3.**
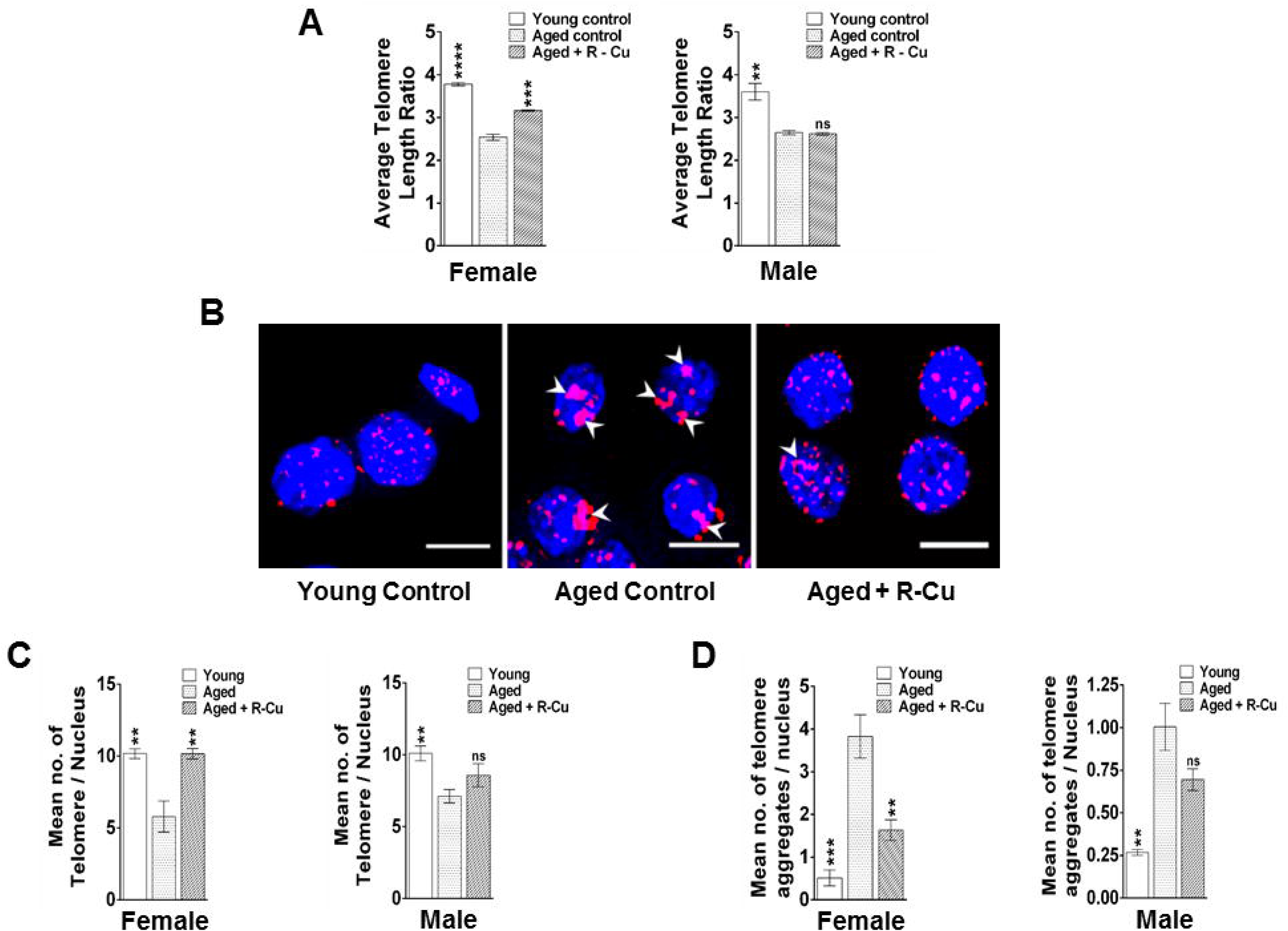
Telomere abnormalities in brain cells of ageing mice and their prevention by treatment with R-Cu. **A.** Estimation of telomere length by quantitative RT-PCR. Bars represent mean ± SEM values. N= 4 animals of both sexes, except in R-Cu treated female mice wherein N was =3; **B.** Representative images of telomere FISH (Scale bar = 10 µm). **C.** Histograms representing number of telomeres per nucleus as estimated by SPOTSCAN software. Mean number of fluorescent telomere signals per nucleus from five randomly chosen fields (∼ 500 nuclei) is represented in the histograms. Bars represent mean ± SEM values. N=4 animals each group of both sexes; **D.** Histograms representing telomere aggregates per nucleus (marked by arrow heads in B). Mean number of fluorescent telomere aggregates per nucleus from five randomly chosen fields (∼ 500 nuclei) is represented in the histograms. Bars represent mean ± SEM values. N=4 animals each group of both sexes. In A, C and D values in young controls and R-Cu treated ageing mice were compared with those of ageing controls, and statistical analysis was performed by two-tailed Student’s t test. ** p < 0.01; *** p < 0.001; **** p < 0.0001.

### R-Cu reduces amyloid deposition in ageing brain and restores of BDNF levels in serum

Increased amyloid (Aβ) protein deposition in extra-cellular spaces of brain cells is classically associated with Alzheimer’s disease [34, 35]. Confocal microscopy using antibody against β-amyloid detected markedly increased amyloid deposition in the form of amyloid fibrils in ageing mice. The latter was remarkably reduced following one year of R-Cu treatment (Fig.4A, left hand panel). Quantification of MFI confirmed the marked increase in extracellular β-amyloid deposition in ageing mice brain (p<0.0001 and p<0.01 in female and male mice, respectively). One year of R-Cu treatment resulted in significant reduction in extra-cellular amyloid in mice of both sexes (p<0.01 and p<0.5 in female and male mice, respectively) (Fig.4A, right hand panel).

**Figure 4.**
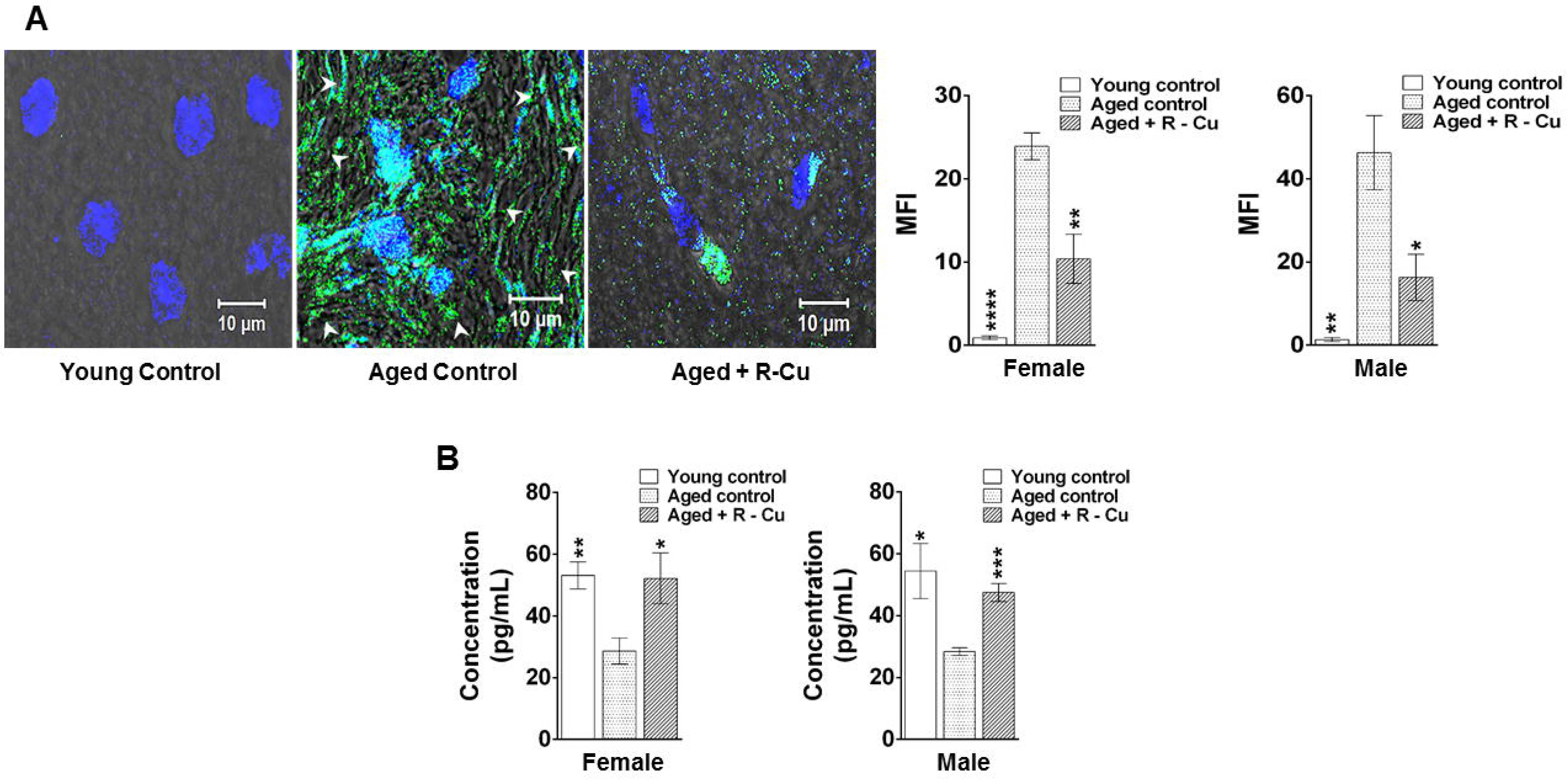
Increased β-amyloid deposition in ageing brain and reduced BDNF in serum, both of which are reversed by treatment with R-Cu. **A.** Representative confocal images of β- amyloid deposition in brain as detected by IF (left hand panel). Quantitative histograms representing mean MFI values (± SEM) of β- amyloid fluorescence (right hand panel). For each slide, 1000 cells were analysed. N=4 animals in each group of both sexes. **B.** Quantitative histograms representing mean serum BDNF levels (± SEM). N=4 animals in each group of both sexes except, young control female (N=3), ageing R-Cu treated female (N=3), and ageing control male mice (N =3). In A and B, levels in young controls and R-Cu treated ageing animals were compared with those of ageing controls, and statistical analysis was performed by two-tailed Student’s t test. * p< 0.05; ** p < 0.01; *** p < 0.001; **** p < 0.0001.

Brain derived neurotrophic factor (BDNF), which plays an important role in neuronal survival and growth [36], was greatly reduced in sera of ageing mice of both sexes as measured by ELISA (p<0.01 and p<0.05 in female and male mice, respectively) (Fig.4B). R-Cu treatment for one year resorted serum BDNF levels nearly to those seen in young mice in both sexes (p<0.05 and p<0.001 in female and male mice, respectively) (Fig.4B).

### R-Cu prevents DNA damage, apoptosis and inflammation in ageing brain cells

We next examined several other hallmarks of ageing viz. DNA damage, apoptosis and inflammation in brain cells [37–39]. DNA damage was examined using phosphorylation of H2AX as a marker of dsDNA breaks [40]. γ-H2AX levels were markedly increased in ageing mice (p<0.001 and p<0.0001 in female and male mice, respectively). R-Cu treatment reduced γ-H2AX levels (p<0.01 and p<0.05 in female and male mice, respectively) (Fig.5A, left & right hand panels).

**Figure 5.**
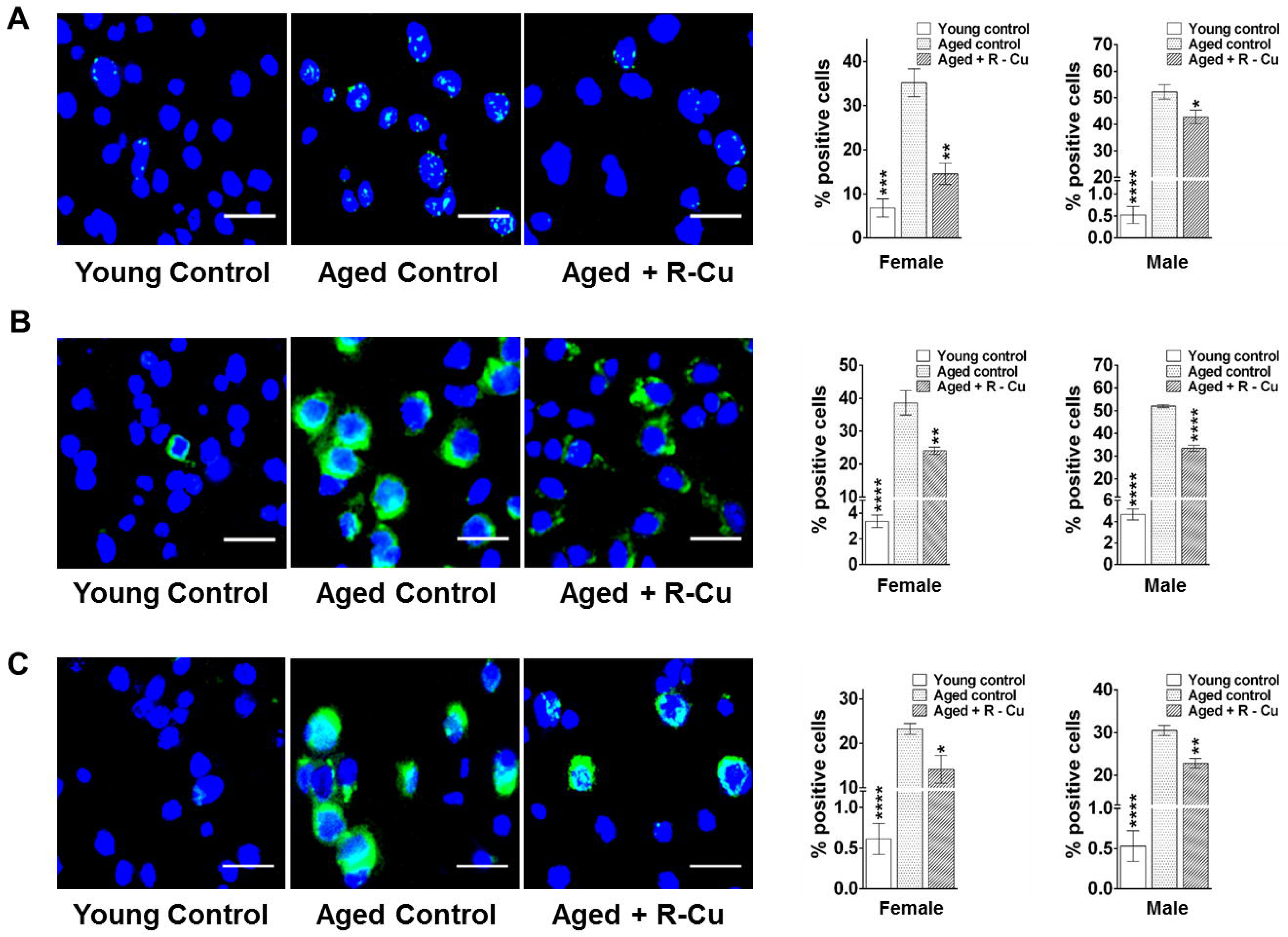
DNA damage, apoptosis and Inflammation in brain cells of ageing mice and their prevention by treatment with R-Cu. A. Representative images of γH2AX expression (Scale bar = 10 µm) (left hand panel). Quantitative histograms (right hand panel). For each slide 1000 cells were analysed and percent cells positive for γH2AX were calculated. Bars represent mean ± SEM values. N=4 animals in each group of both sexes. **B.** Representative images of active caspase 3 expression (Scale bar = 10 µm) (left hand panel). Quantitative histograms (right hand panel). For each slide 1000 cells were analysed and percent cells positive for caspase-3 were calculated. Bars represent mean ± SEM values. N=4 animals in each group of both sexes. **C.** Representative images of NF-kB expression (Scale bar = 10 µm) (left hand panel). Quantitative histograms (right hand panel). For each slide 1000 cells were analysed and percent cells positive for NF-kB were calculated. Bars represent mean ± SEM values. N=4 animals in each group of both sexes. For A, B and C, levels in young controls and R-Cu treated ageing animals were compared with those of ageing controls, and statistical analysis was performed by two-tailed Student’s t test. * p< 0.05; ** p < 0.01; *** p < 0.001; **** p < 0.0001.

We next examined active caspase-3 levels, which is known to be a marker of mitochondria mediated apoptosis [41]. Highly significant increase in apoptosis in ageing brain cells was seen in both sexes (p< 0.0001). R-Cu treatment significantly reduced the degree of apoptosis in mice of both sexes (p<0.01 and p< 0.0001 in female and male mice, respectively) (Fig.5B, left & right hand panels).

Inflammation is a cardinal hallmark of ageing [12], and we assessed the expression of the transcription factor NF-kB in brain cells. The latter was significantly elevated in ageing mice of both sexes (p< 0.0001). R-Cu treatment significantly reduced NF-kB levels in both sexes (p< 0.05 and p< 0.01 in female and male mice, respectively) (Fig.5C, left & right hand panels).

### R-Cu prevents senescence in ageing brain cells

Senescence is the hallmark of biological ageing characterised by gradual deterioration of cellular functions [42]. We observed persistence of numerous co- localizing signals of γ-H2AX with those of telomeres (DNA-SCARS) - a classical hallmark of senescence [43] in ageing mice of both sexes (p< 0.001). Quantification of co-localising signals revealed a marked reduction following R-Cu treatment (p<0.01 in both sexes) (Fig.6A, upper & lower panels).

**Figure 6.**
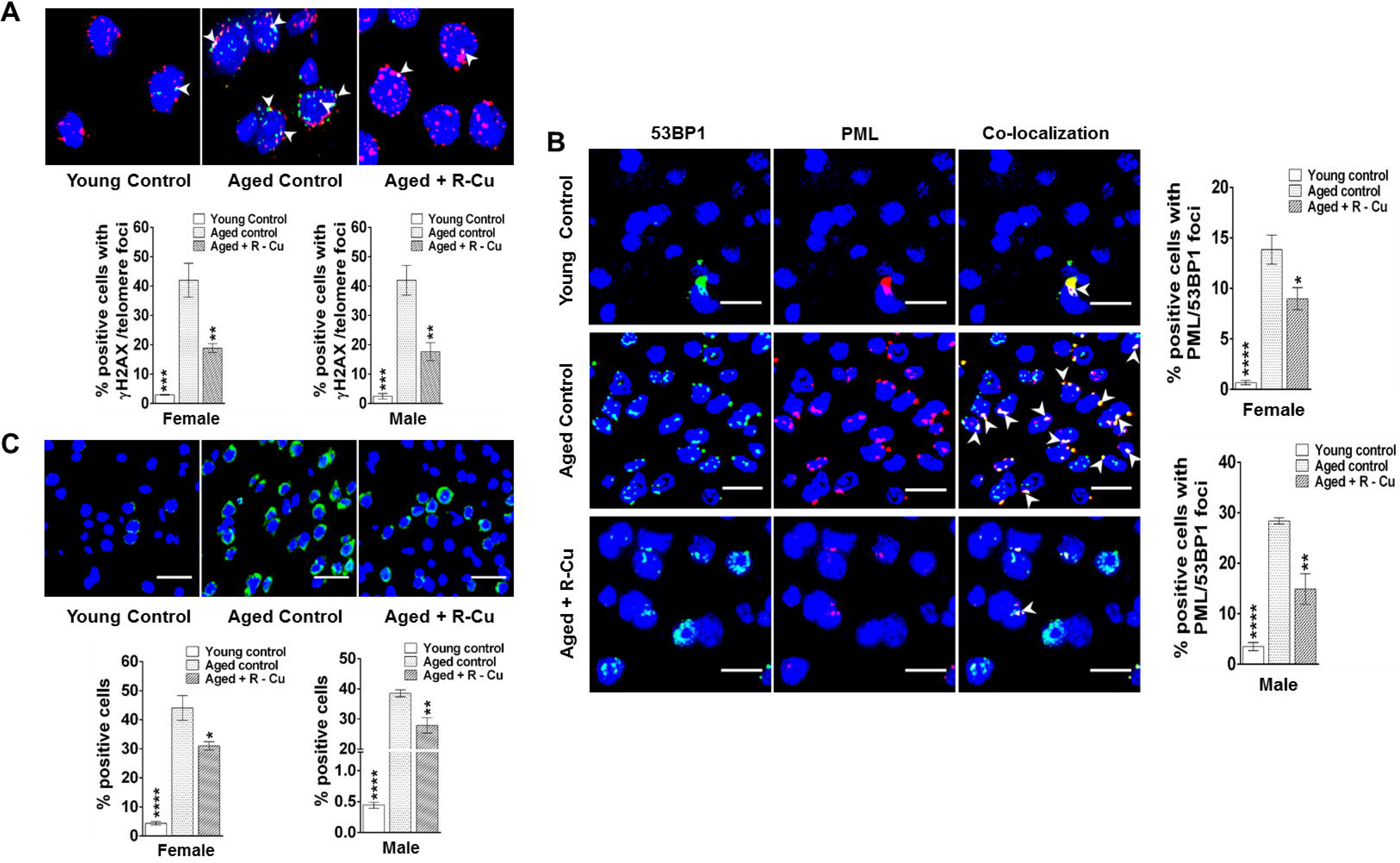
Activation of hallmarks of senescence in brain cells of ageing mice and their prevention by treatment with R-Cu. A. Representative immuno-FISH images showing co-localization of fluorescent signals of γ-H2AX and telomeres (upper panel) (Scale bar = 10 µm). Histograms of quantitative estimation of co- localised signals (lower panel). For each slide 500 nuclei were analysed and percent cells showing co-localisation of γH2AX and telomere signals were calculated. Bars represent mean ± SEM values. N=4 animals in each group of both sexes. **B.** Representative IF images showing co-localization of fluorescent signals of 53BP1 and PML (left hand panel) (Scale bar = 10 µm). Histograms of quantitative estimation of co-localised signals (right hand panel). For each slide 500 nuclei were analysed and percent cells showing co-localisation of 53BP1and PML signals were calculated. Bars represent mean ± SEM values. N=4 animals in each group of both sexes. **C.** Representative images of p16 expression (upper panel) (Scale bar = 10 µm) and quantitative histograms (lower panel). For each slide 1000 cells were analysed and percent cells positive for p16 was calculated. Bars represent mean ± SEM values. N=4 animals in each group of both sexes. In A, B and C, values in young controls and R-Cu treated ageing animals were compared with those of ageing controls, and statistical analysis was performed by two-tailed Student’s t test. * p< 0.05; ** p < 0.01; *** p < 0.001; **** p < 0.0001.

Double strand DNA breaks form a storage hub for several heterochromatin-binding proteins in the form of senescence-associated heterochromatic foci (SAHF) [44]. One such heterochromatin binding protein is pro-myelocytic leukemia-nuclear bodies (PML-NBs) which has been shown to be significantly correlated with DNA damage associated senescence in ageing mice [45, 46]. We show that the number of co- localizing signals of 53BP1 and PML were markedly increased in ageing mice of both sexes (p<0.0001). Co-localising signals were significantly reduced following R- Cu treatment for 1 year (p< 0.05 and p< 0.01 in female and male mice, respectively) (Fig.6B, left & right hand panels).

Another marker of senescence that we examined was p16 ^INK4a^ [47], which was elevated in ageing mice of both sexes (p< 0.0001 for both sexes). R-Cu treatment significantly reduced levels of p16 ^INK4a^ (p< 0.05 and p<0.01 in female and male mice, respectively) (Fig.6C, upper & lower panels).

### R-Cu prevents aneuploidy in ageing brain cells

Telomere loss resulting in fusion of chromosomes in ageing mice can cause aneuploidy resulting in abnormal number of chromosomes [48]. We examined the degree of aneuploidy in brain cells with respect to chromosome no 7 and 16, and observed a ∼ 15 fold increase in aneuploidy in ageing mice with respect to both chromosomes (p<0.0001 for both chromosomes in both sexes) (Fig.7) left & right hand panels). R-Cu treatment markedly reduced aneuploidy with respect to both chromosomes in both sexes (p<0.001)) (Fig.7, left & right hand panels).

**Figure 7.**
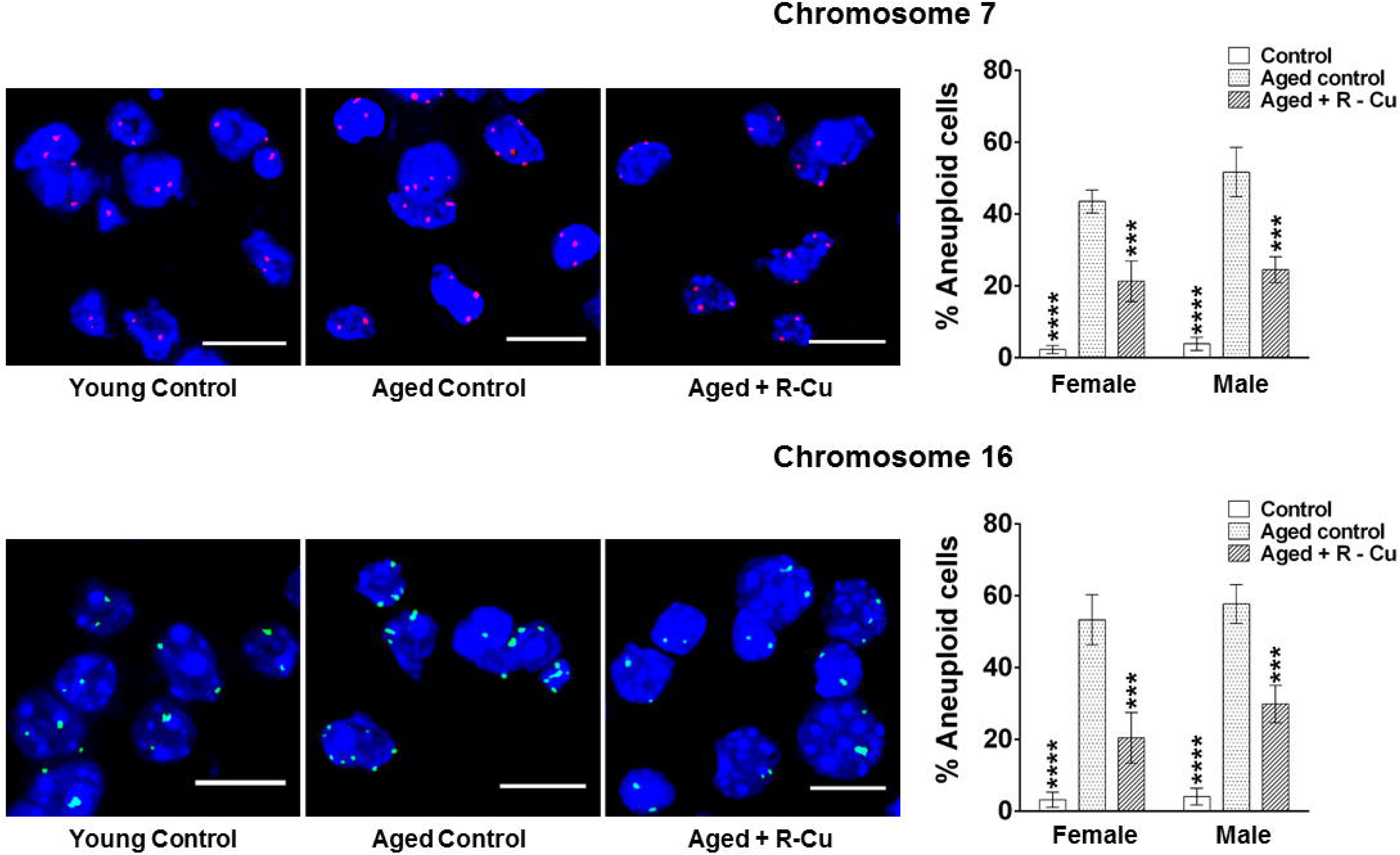
Development of aneuploidy in brain cells of ageing mice and its prevention by treatment with R-Cu. Representative images of aneuploidy of chromosome 7 and chromosome 16 in brain cells (left hand panels) (Scale bar = 10 µm). Quantitative histograms representing percent aneuploid cells (right hand panels). Number of fluorescent signals per nucleus was counted, and signals less than 2N or more than 2N in a nucleus were taken as evidence of aneuploidy. Five fields (∼ 500 nuclei) were analysed and average number of signals per nucleus was calculated. Bars represent mean ± SEM values. N=4 animals in each group of both sexes. Values in young controls and R-Cu treated ageing animals were compared with those of ageing controls, and statistical analysis was performed by two-tailed Student’s t test. *** p < 0.001; **** p < 0.0001.

### R-Cu prevents mitochondrial dysfunction in ageing brain cells

Mitochondrial DNA integrity and functionality decrease with age resulting in accumulation of oxidative damage caused by reactive oxygen species (ROS) [49]. We studied mitochondrial dysfunction by analysing the expression of TOM20, a nuclear-encoded subunit of the mitochondrial translocation complex which imports other nuclear-encoded proteins. Its overexpression is reported to promote neurodegeneration [50]. Our results revealed overexpression of TOM20 on mitochondrial membrane leading to increase in total mitochondrial volume in ageing brain cells when compared to young controls (p< 0.001 in both sexes). R-Cu treatment significantly restored mitochondrial volume (p< 0.05 and p<0.01 in female and male mice, respectively) (Fig.8, upper & lower panels).

**Figure 8.**
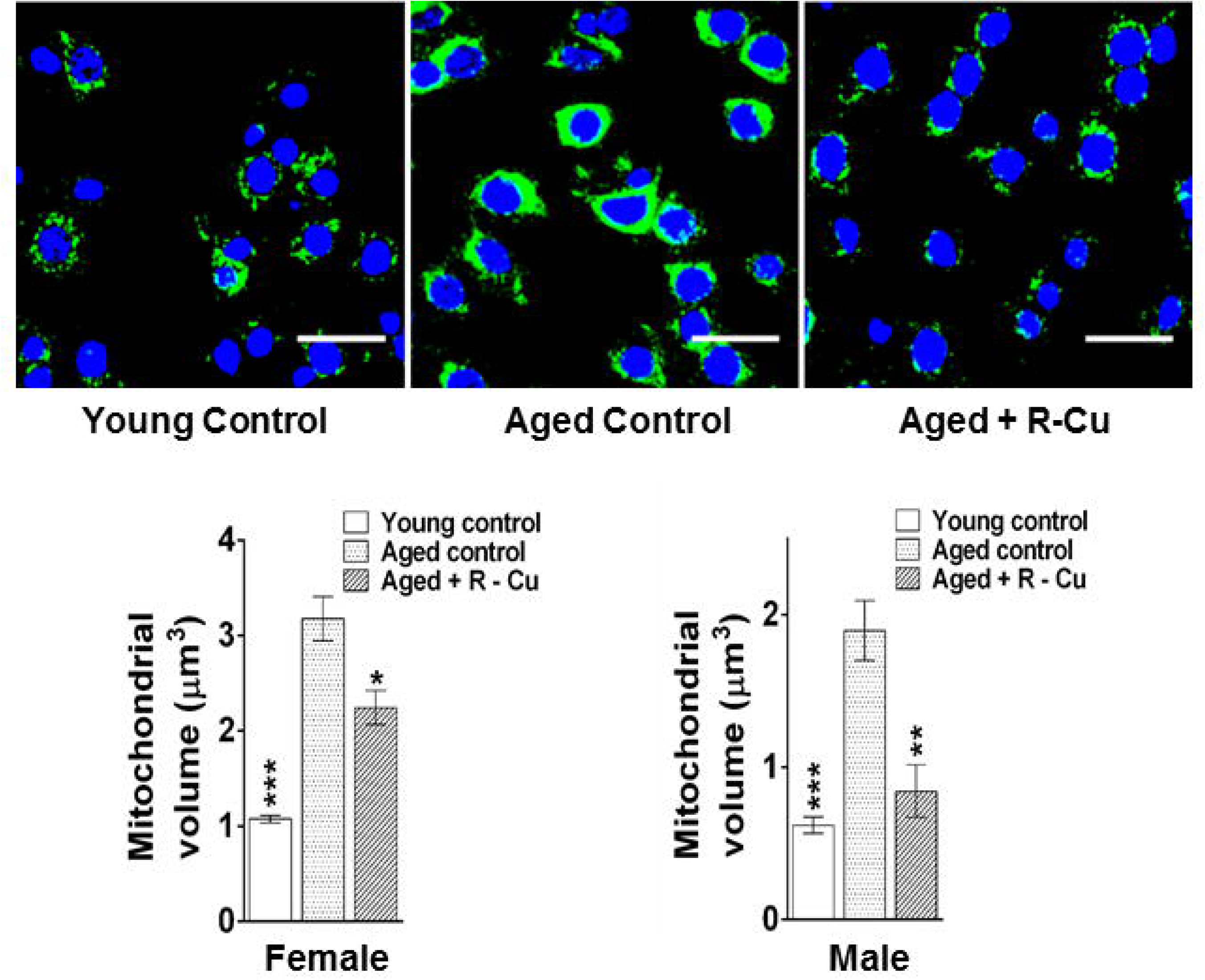
Increased mitochondrial dysfunction in ageing mice and its restoration by treatment with R-Cu. Representative IF images showing expression of TOM20 (upper panel) (Scale bar = 10 µm). Quantitative histograms representing mitochondrial volume changes (lower panel). Five fields (∼ 2000 mitochondria) were analysed and average volumetric change was estimated using IMARIS software. Bars represent mean ± SEM values. N=4 animals in each group of both sexes. Values in young controls and R-Cu treated ageing animals were compared with those of ageing controls, and statistical analysis was performed by two-tailed Student’s t test. * p< 0.05; ** p < 0.01; *** p < 0.001.

### R-Cu prevents systemic metabolic dysfunction in ageing mice

Metabolic ageing involves dysregulation of physiological processes leading to insulin resistance and lipid accumulation brought about by low grade chronic inflammation [51, 52]. As anticipated serum glucose levels was significantly elevated in ageing mice [53] (p< 0.01 for both sexes), which was reduced to levels seen in young controls following one year treatment with R-Cu (p< 0.01 for both sexes) (Fig.9A). Serum cholesterol was significantly elevated in ageing female mice (p<0.05) but not in male mice (Fig.9B). Nonetheless, R-Cu treatment reduced serum cholesterol levels well below that observed in young control mice of both sexes (p< 0.001 in both sexes). CRP was highly significantly elevated in ageing mice of both sexes (p<0.0001) and was reduced by R-Cu treatment (p<0.05 in both sexes) (Fig.9C).

**Figure 9.**
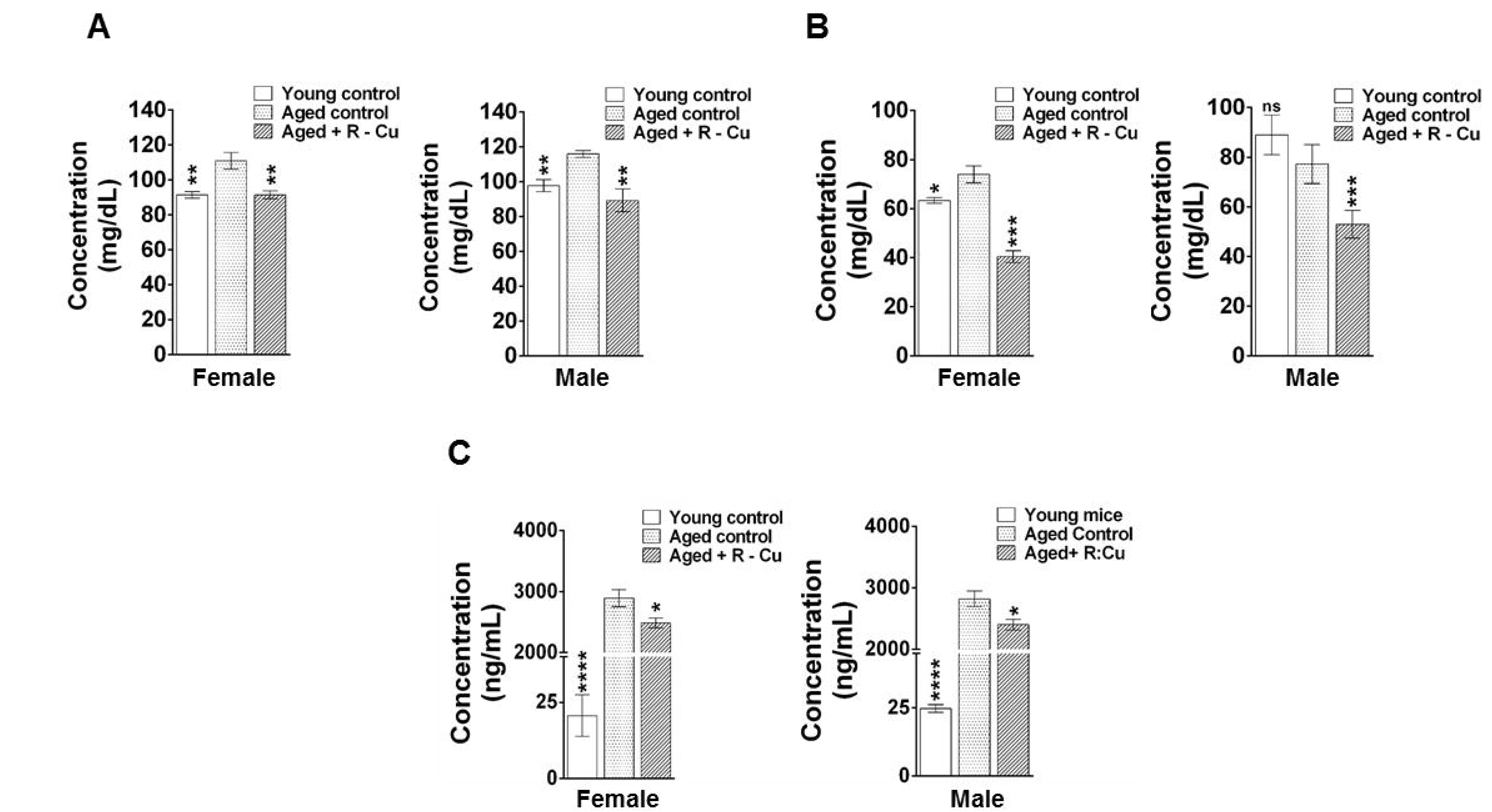
Increased metabolic dysfunction in ageing mice and its prevention by treatment with R-Cu. A, B and C. represent histograms of levels of serum glucose, cholesterol and CRP, respectively. Bars represent mean ± SEM values. N=4 animals in each group of both sexes except in cholesterol female young control (N=3) and cholesterol male young control (N=2). Levels in young controls and R-Cu treated ageing animals were compared with those of ageing controls, and statistical analysis was performed by two-tailed Student’s t test. * p< 0.05; ** p < 0.01; *** p < 0.001; **** p < 0.0001.

## Discussion

ROS are short lived molecular species containing an unpaired electron which makes them highly reactive as they search for another electron to pair with, and in the process can damage biomolecules such as DNA, proteins and lipids [54]. ROS induced oxidative stress is known to have several harmful effects on host cells [55]. However, we have reported that, paradoxically, when ROS is artificially generated outside the cell in the extracellular spaces of the body, they can have wide ranging therapeutic effects [18-20, 26, 27]. Admixing R with Cu leads to generation of oxygen radicals by virtue of the ability of R to reduce Cu (II) to Cu (I) [23, 25]. Oxygen radicals that are generated in the stomach upon oral administration of R-Cu are readily absorbed to have systemic effects in the form of deactivation / eradication of extracellular cfChPs. The latter have multiple deleterious effects in the host. For example, cfChPs can readily enter into the healthy cells to damage their DNA, activate inflammatory cytokines and promote apoptosis via the mitochondrial pathway [13, 14]. Given that 1X10^9^-1X10^12^ cells die in the body every day [56, 57], we have hypothesised that repeated and lifelong assault on healthy cells by cfChPs derived from the dying cells might be the underlying cause of ageing [15, 16]. In support of this hypothesis we show in this article that prolonged oral administration of R-Cu to ageing mice down-regulated multiple biological hallmarks of ageing and neurodegeneration. An illustrated summary of the mechanisms by which R-Cu generated oxygen radicals eradicate cfChPs from brain micro-environment to down- regulate biological hallmarks of ageing is provided in Fig.10. Our results suggest that R-Cu may qualify as an ideal anti-ageing agent since it has the potential to simultaneously retard or delay the many conditions that are associated with ageing [2]. To be globally applicable, an ideal anti-ageing agent should also be inexpensive and non-toxic – the two criteria that are also met by R-Cu. The latter can be easily administered orally, and both R and Cu are already approved for human use.

**Figure 10.**
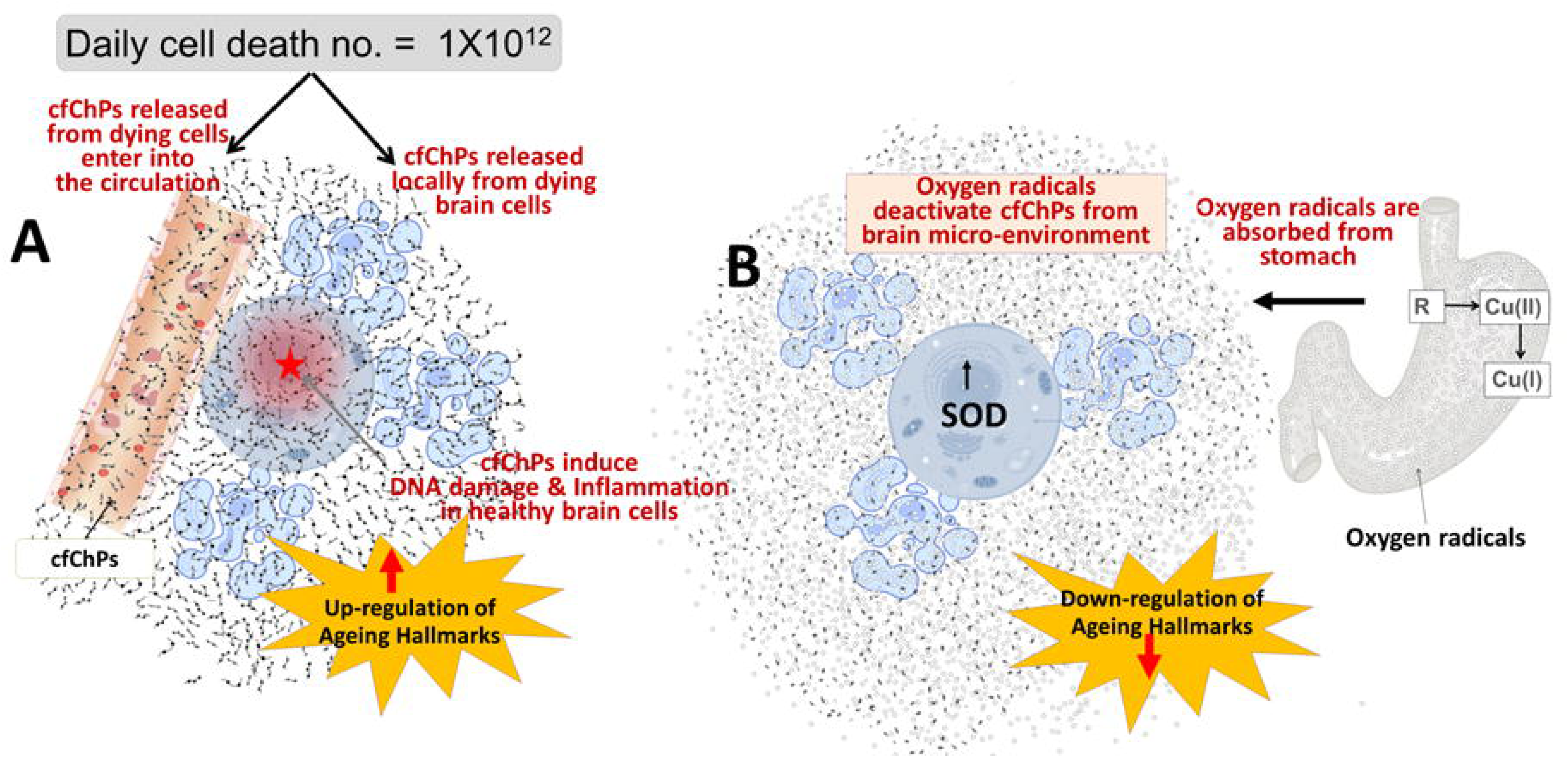
Graphical illustration of mechanisms involved in oxygen radical induced down-regulation of biological hallmarks in ageing in brain cells. **A**. cfChPs that are extruded from circulation, or those that are released locally from dying brain cells, are readily internalised by healthy brain cells, wherein they activate multiple biological hallmarks of ageing. **B**. Oxygen radicals are generated in the stomach upon oral administration of R-Cu which are readily absorbed leading to systemic effects in the form of deactivation / eradication of cfChPs in the brain micro- environment. Deactivation / eradication of cfChPs leads to down-regulation of biological hallmarks of ageing in brain cells. Oxygen radicals also enter into the healthy brain cells; but their entry leads to activation of the anti-oxidant enzyme superoxide dismutase (SOD), which detoxifies and eliminates the offending agents, thereby protecting the genomic DNA.

The mechanism(s) by which R-Cu down-regulates the multiple biological hallmarks of ageing and neurodegeneration needs elaboration. Reversal of telomere shortening by R-Cu may suggest that telomere shortening could be a consequence of dsDNA breaks inflicted by cfChPs which shear off telomere ends causing them to shorten. This may also help to explain our detection of persistent γ-H2AX signals in telomere regions of brain cells, which is considered to be a signature of senescence [43]. The bare chromosomal ends may fuse with each other to lead to chromosomal instability and aneuploidy [48], as was detected in our study. With respect to mitochondria, we have recently reported that internalised cfChPs can inflict physical damage to mitochondria, and that one of the indicators of mitochondrial damage detected in that study was up-regulation of TOM20 [58]. Our finding in the current study that TOM20 expression in ageing mice could be reversed by R-Cu is consistent with the possibility that mitochondrial damage in ageing mice is induced by cfChPs. However, reduction in amyloid plaque formation following prolonged R- Cu treatment would suggest a yet unknown role of cfChPs which calls for further research. Likewise, the mechanism(s) by which R-Cu reduced metabolic dysfunction in ageing mice leading to reduction in serum levels of glucose, cholesterol and CRP remains unknown at present. Taken together, it can be concluded that cfChPs have pleiotropic effects with wide ranging implications in ageing and neurodegeneration which remain open to future research.

We did not detect any evidence of damage to genomic DNA of brain cells that could be attributed to oxygen radicals generated following one year of R-Cu treatment. The markedly up-regulated antioxidant enzyme SOD apparently neutralised the invading oxygen radicals and prevented damage to cellular genomic DNA (vide Fig.1). This was further substantiated by our finding of a reduction in γ-H2AX signals in the post R-Cu treated mice (vide Fig.5A). Overall, we observed no adverse effects in mice which had been administered R-Cu for a period of one year. This suggested that genomes of all cells of the body were similarly protected from the potentially damaging effects of oxygen radicals by up-regulated anti-oxidant enzymes.

We demonstrate for the first time that cfChPs derived from dying brain cells are abundantly present in the extracellular spaces of the ageing brain, and that they were virtually eliminated following prolonged treatment with R-Cu. The fact that elimination of cfChPs was associated with down-regulation of multiple biological hallmarks of ageing and neurodegeneration makes a strong case for a direct role of cfChPs in the aetiology of these conditions. We propose that cfChPs released from dying brain cells initiate a vicious cycle of more DNA damage, apoptosis and inflammation, setting in motion a low grade and unrelenting “cytokine storm” [59]. We propose that the latter, together with the other yet unknown harmful pleiotropic effects of cfChPs, are the underlying processes that define ageing. Our results suggest that these harmful effects can be prevented by deactivation / eradication of the offending cfChPs via the medium of oxygen radicals. We propose that oral administration of a combination of small quantities of R and Cu holds the promise of being an effective anti-ageing therapeutic combination. Whether R-Cu will be effective in retarding ageing and neurodegeneration in humans will have to await clinical trials in appropriate populations. It is to be noted, however, that our early results have shown that R-Cu is therapeutically effective in humans, albeit in context of other pathological situations [26, 27].

## Acknowledgement

We sincerely thank Dr. Snehal Shabrish for preparing the info graphic and Mr. Roshan Shaikh and Mr. Ashish Pawar for their help in preparing the manuscript.

## Funding

This study was supported by the Department of Atomic Energy, Government of India, through its grant CTCTMC to Tata Memorial Centre awarded to IM.

## Declaration of competing interests

The authors declare no competing interests.

## Author contributions

K.P., G.V.R., J.D., S.M., V.J., A.S., B.R., H.T., D.K. and S.C. conducted the experiment. I.M., K.P. and G.V.R supervised the experiments. K.P. and G.V.R conducted data analysis. I.M. conceptualized the project, was responsible for overall supervision and procured funding. I.M., K.P. and G.V.R. wrote the paper. I.M. approved the final manuscript.

## Notes

### Competing Interest Statement

The authors have declared no competing interest.

