## Supplementary Table for "A pro-oxidant combination of resveratrol and copper down-regulates multiple biological hallmarks of ageing and neurodegeneration"

### Supplementary Table 1

#### Description of Kits

| Sr. No. | Kits | Catalogue No. | Company / Vendor |
|---|---|---|---|
| 1. | QIAamp® DNA FFPE Tissue Kit (50) | 56404 | Qiagen, Hilden, Germany |
| 2. | Total BDNF Quantikine ELISA Kit | DBNT00 | R&D Systems, MN, USA |
| 3. | Mouse C-Reactive Protein /CRP Quantikine ELISA Kit | MCRP00 | R&D Systems, MN, USA |

#### Description of Antibodies

| Sr. No. | Antibody / Probes | Catalogue No. | Company / Vendor |
|---|---|---|---|
| 1. | Histone H4 IgG | Custom synthesized | Bioklone Biotech Pvt Ltd, Chennai, India |
| 2. | Anti-DNA antibody | NB110-89473 | Novus Biologicals, Colorado, USA |
| 3. | Anti- γ H2AX (Phospho S139) antibody | ab26350 | Abcam, Cambridge, UK |
| 4. | Anti NF-kB p65 antibody | ab32536 | Abcam, Cambridge, UK |
| 5. | Anti-53BP1 antibody (EPR2172(2)) | ab175933 | Abcam, Cambridge, UK |
| 6. | Anti-CDKN2A/p16INK4a antibody (2D9A12) | ab54210 | Abcam, Cambridge, UK |
| 7. | Anti-Cleaved Caspase-3 antibody | ab2302 | Abcam, Cambridge, UK |
| 8. | Anti-TOMM20 antibody (EPR15581-54) | ab186735 | Abcam, Cambridge, UK |
| 9. | Purified anti-β-Amyloid, 1-42 Antibody | 805501 | BioLegend, California, USA |
| 10. | Anti-Superoxide Dismutase 1 antibody | ab13498 | Abcam, Cambridge, UK |
| 11. | Anti-PML antibody | P6746 | Merck-Millipore Sigma, Germany |
| 12. | Goat Anti-Rabbit IgG (H+L) FITC conjugate secondary antibody | AP307F | Merck-Millipore Sigma, Germany |
| 13. | Goat Anti-Mouse IgG H&L (FITC) pre-adsorbed secondary antibody | ab7064 | Abcam, Cambridge, UK |
| 14. | Rabbit Anti-Goat IgG H&L (FITC) secondary Antibody | ab6737 | Abcam, Cambridge, UK |
| 15. | Goat anti-mouse IgG H& L (FITC) secondary Antibody | ab6785 | Abcam, Cambridge, UK |
| 16. | PNA Telomere FISH probe ( Tel-G-Cy3) | F1006 | Panagene, South Korea |
| 17. | Single chromosome paint probes Chromosome 7 red (Ready to use) | Custom synthesized | Applied Spectral Imaging, Israel |
| 18. | Single chromosome paint probes Chromosome 16 green | Custom synthesized | Applied Spectral Imaging, Israel |
